# A literature mining method to judge whether there are uncertainties in empirical-dependent antineoplastic drug distribution in specific clinical scenarios

**DOI:** 10.1101/842401

**Authors:** Xiaoyang Ji, Zhendong Feng, Qiangzu Zhang, Zhonghai Zhang, Yanhui Fan, Renhua Na, Gang Niu

**Affiliations:** Key Laboratory of Animal Genetics, Breeding and Reproduction of Inner Mongolia Autonomous Region College of Animal Science, Inner Mongolia Agricultural University, Hohhot 010018, China; Phil Rivers Technology, Beijing, China; State Key Laboratory of Computer Architecture, Institute of Computing Technology, Chinese Academy of Sciences, Beijing, China

**Keywords:** Literature mining method, Antineoplastic drug, Different cancer types, Linear regression analysis, Antineoplastic drug-gene association matrix

## Abstract

Cancer clinical practice guidelines recommend different treatment options for different cancer types and are mainly developed by clinicians. In theory, those recommendation schemes that are supported by scientific research should provide better efficacy for patients. However, in actual clinical practice: “Is the choice of a specific antineoplastic drug for a specific cancer supported by the results of molecular biology mechanisms or based on the subjective experience of the clinician?” Answering this question is of significant importance for guiding clinical practice, but there is currently no operational method to provide objective judgment in specific cases. This paper describes a literature mining method that collates information from specific antineoplastic drug-related literature to establish an antineoplastic drug-gene association matrix for global or specific cancer scenarios, and further establishes a standard model and scenario models. Based on the parameters of these models, we constructed a linear regression analysis method to evaluate whether the models in different scenarios deviated from a random distribution. Finally, we determined the possible efficacy of an antineoplastic drug in different cancer types, which was validated by the Genomics of Drug Sensitivity in Cancer (GDSC) database. Using our mining method, we tested 18 antineoplastic drugs in 16 cancer types. We found that cisplatin used in ovarian cancer was more efficacious and may benefit patients more than when used in breast cancer, which provides a new paradigm for rational knowledge-driven drug distribution patterns in clinical practice.

## Introduction

More and more in-depth studies of antineoplastic drugs have revealed that the same antineoplastic drug can have drastically different effectiveness in tumors originating from different tissues. Excluding a limited number of potential immune pharmaceuticals that have broad-spectrum activity against a wide range of cancer types ^[1, 2]^, the majority of approved chemotherapeutic drugs and targeted drugs, as well as their combinations, generally target specific pathological types and specific cancer types for which they were primarily intended. Clinicians in different cancer research fields often recommend different treatment options for specific cancer types based on clinical practice guidelines that are primarily aggregated by experience. Given that clinical trials are conducted on humans, researcher must fully consider the complexities of these studies, including the type of trial, subjects, controls, sample sizes, main outcomes, as well as their implementation. In addition, clinical trials are limited by time and cost, and ethical issues, as well as subject to relevant methodological quality assessments. Therefore, most expert consensus statements in clinical practice guidelines are often based on 1) the urgency of the clinical needs, 2) the small sample size and data size of clinical trials, 3) the clinical experience of experts, and 4) the collation and analysis of some published literature results in related fields, which often leads to uncertain therapeutic effects after the implementation of clinical drug regimens. For example, sorafenib is the first-line treatment for advanced hepatocellular carcinoma (HCC), but it is not effective in Chinese HCC patients. More than 90% of HCC cases in China are caused by hepatitis B virus (HBV) infection^[3]^, and sorafenib has limited effect in HCC caused by HBV^[4, 5]^. This shows that the current application of antineoplastic drugs in different clinical treatments for cancer still mainly depend on experience, with considerable uncertainty.

In the past decade, there has been an increasing number of scientific studies on the application and mechanisms of specific antineoplastic drug in different cancer scenarios. This research can be accessed in various literature databases. For example, we searched the PubMed database on October 9, 2019 and retrieved 72,820 papers with ‘cisplatin’ as the keyword and 6,528 papers with ‘erlotinib’ as the keyword ^[6]^. With rapid developments in biomedical research, the amount of scientific literature has increased rapidly, and it is becoming increasingly difficult for human researchers to assimilate all the knowledge related to a subject. Therefore, the research models adopted for antineoplastic drugs by researchers are more likely to follow a causal relationship hypothesis proposed by researchers based on incomplete information, and such incomplete research might form the description of the pharmacological mechanisms. Subsequently, this incomplete research mechanism may be used to refine search key terms to screen references supporting this deficient research hypothesis in the literature databases, and finally form a description of the antineoplastic drug mechanism guided by subjective experience. This incomplete grasp of the information leads to “subjective” operation in the use of prior knowledge by a single human researcher, which strengthens the empirical judgment of the causal relationship hypothesis.

This study adopted a gene-based literature mining method to establish a standard model of an antineoplastic drug and different scenario models of the antineoplastic drug in various cancer scenarios by constructing a gene-antineoplastic drug association matrix displayed in a standardized interface. We can initially answer the following question using these models: “Is the application of a specific antineoplastic drug in an actual clinical scenario of cancer based on a molecular biology mechanism derived from solid scientific evidence (rational basis), or is the judgment based on human subjective experience (empirical basis)?” Experience-based treatment outcomes tend to use drugs randomly, whereas scientific research results can lead to better clinical outcomes. Therefore, we have established a mathematical model for judging whether medication utilization is experience-based or evidence-based in a specific scenario, which can then be validated using the Genomics of Drug Sensitivity in Cancer (GDSC) database. This method not only evaluates the applicability of antineoplastic drugs in different cancer types, but also automatically delineates its biological mechanisms, helping researchers better understand the efficacy of the antineoplastic drug for various cancer types, as well as the possible causes of tolerance.

## Methods

### Construction of data interface

We used PubMed to provide the biological literature for the text mining. The schematic representation of the overall study architecture is shown in Fig 1 and can be summarized in the following steps. Step 1, information retrieval. PubMed is searched and relevant information is downloaded and used to build the subject dictionary (SD) and public health dictionary (PD). Step 2, identify gene entities accurately related to the subject. Step 3, identify all biological entities that are accurately related to the subject (disease, drugs, phenotypes, treatments, and other relevant clinical terms). Step 4, calculate and rank the strength of associations between extracted genes and entities. Step 5, establish association matrix between subject-related antineoplastic drug entities and genes by alignment with antineoplastic drug name database. The details of each step are described below and each algorithm consists of custom scripts unless otherwise stated.

**Fig. 1.**
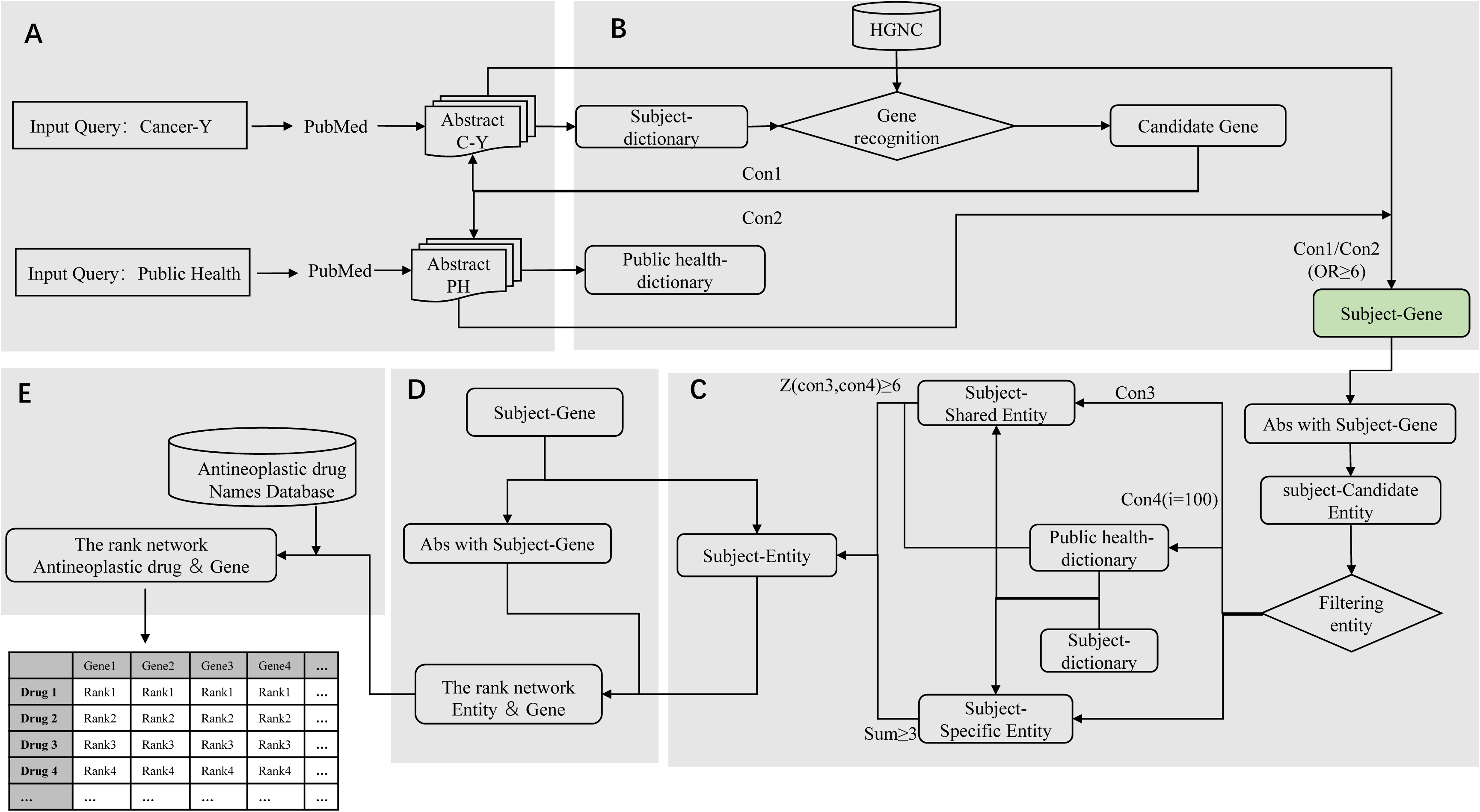
Overview of the components of literature mining. The process of retrieving evidence-based sentences from PubMed abstracts and the basic steps of the literature mining: (A) retrieve information, (B) identify genes with accurate relevance to the subject, (C) identify entities with accurate relevance to the subject, (D) calculate the association strengths between entities and genes, and (E) align with antineoplastic drug name database to establish an association matrix between antineoplastic drug subject-entities and genes.

#### Step 1: Information retrieval

The dataset used in this pipeline uses only PubMed articles. First, PubMed is searched for articles containing the subject keywords, including abstracts, titles, and author/unit information sections. The search results are downloaded in txt format to obtain structured information. Then, the text in the subject abstract set is organized and cleaned, and compiled into the subject dictionary (SD). To enhance the accuracy of the effective entities associated with the keyword, we use a random corpus for comparison. We search for article abstracts containing “public health” as the keyword and compile this abstract set into the public health dictionary (PD), which contains a wide range of proteins, genes, and related biological entities. Meanwhile, we also consider the balance of the amount of information by setting relevant parameters to adjust the amount of text before carrying out the statistical analyses.

#### Step 2: Identify gene entities precisely related to the subject

Biological entity identification is a key step in the literature mining process^[7, 8]^. To ensure functionality of the extracted entity, we first compare the entity from SD with the human official gene symbols in the Hugo Gene Nomenclature Commission (HGNC)^[9]^ database to generate subject candidate genes using standard nomenclature. In addition, the entities in the abstract are capitalized to avoid errors in the identification process. To obtain widely used gene entities that are precisely related to the subject, we search for the subject candidate genes in the SD and the PD, respectively, and count the number of abstracts containing each subject candidate gene in each abstract set, respectively. Finally, we calculate the odds ratio of each subject candidate gene and sort them into a list of precisely related gene entities. Formula (1) is used to calculate the odds ratio of each gene:

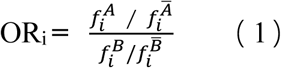

where i is a subject candidate gene, f is the number of abstracts, A is the subject abstract set containing gene i, 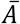 is the subject abstract set that does not contain gene i, B is the public health abstract set containing gene i, and 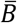 is the public health abstract set that does not contain gene i. The screening criteria for genes precisely related to a subject is 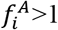, ORi⩾6. As we are focusing on extracting subject-gene associations, we retain only those abstracts that have at least one subject-gene mentioned and define this as the subject gene abstract set (SGA).

#### Step 3: Identify all biological entities that are accurately related to the subject

We first compare the entities in PD and SD to obtain the subject-specific entity dictionary (SPE) containing unique entities in SD and the subject-shared entity dictionary (SHE) containing shared entities between PD and SD. To further improve the accuracy of the recognition rate of the subject-related entities, we compare SGA with the entities in SPE and SHE, respectively. We first perform a comparison screening in SHE. We count the number of abstracts containing each subject-shared entity (HEj) in SGA. Next, the same number of abstracts in SGA are randomly extracted from the public health abstract dataset as the reference abstract dataset and this is repeated 100 times. For each randomly extracted reference abstract dataset, it is compared with the entities in SHE and the number of abstracts containing each subject-shared entity (HEj) is counted. The standard score of each entity (HEj) is then calculated in SGA and in the reference abstract set to obtain the first part of the entity precisely related to the subject. Formula (2) is used to calculate the standard score of each entity:

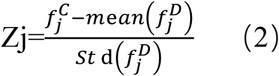

where j is the entity in the SHE, *f* is the number of abstracts, C is the SGA, and D is the randomly extracted reference abstract set from public health abstract set. The screening criteria for the first part of the candidate subject entity is Zj⩾6.

We next perform an alignment screening in SPE. We count the number of abstracts containing each subject-specific entity (PEk) in SGA. If the number of abstracts containing PEk is more than 3 in SGA, then this is designated as the second part of the subject-related entity. A consensus of ≥ 3 has been decided by the authors, with the convention that < 3 articles published may be a random co-occurrence or without any unidirectional scientific evidence. Hence, the articles with <3 may not be of significance. Finally, the two parts are merged to obtain the subject-related entity. In addition, some entities have singular and plural noun forms, and synonyms with multiple forms in the context of the abstract. Therefore, we number the subject-related entity and automatically combine the nouns with plural forms and the homologous words with adjectives and adverb roots into the same entity and assign this same number.

#### Step 4: Calculate and rank the strength of associations between extracted genes and entities

We first need to define the association between gene and entity. If “n” entities and “m” genes co-occur in any literature abstract, the algorithm will automatically extract all “n” and “m” possible entity-gene pairs. The extraction algorithm is probabilistic and does not consider the syntactic relationships between entities and gene entities independently in sentences. The strength of their association is then analyzed by calculating the relationship distance between the subject gene and the entity. For each gene Gi that is precisely related to the subject, the abstract set containing the gene Gi and the abstract set not containing the gene Gi are extracted from SGA. Then, we count the average number of abstracts containing the subject-related entity Ej in the abstract set AGi containing the gene Gi; and count the average number of abstracts containing the subject-related entity Ej in the abstract set NAGi not containing gene Gi according to co-occurrence. We then calculate the difference in the average number of abstracts containing entity Ej between the abstract set AGi and NAGi. We thus obtain the first relationship distance RD_1ij_ between subject gene Gi and subject entity Ei. To obtain more meaningful results and filter possible false positives, we define the significance of the relationship distance by standard scores. First, we mark the subject gene abstracts in the SGA and then rank the abstracts markers randomly, so that the corresponding relationship between the abstract marker and the abstract content is randomly changed to generate 100 random matrices. Then, we separately count the average number of abstracts containing the entity Ej in the abstract set SAGi containing gene Gi; and count the average number of abstracts containing the entity Ej in the abstract set SNAGi not containing gene Gi from the random subject gene abstract set generated by each random cycle. We then calculate the difference of the average number of abstracts containing entity Ej between the abstract set SAGi and SNAGi. We thus obtain the second relationship distance RD_2ij_ between subject gene Gi and subject entity Ej. Next, we determine the relationship distance between subject gene Gi and subject entity Ej by calculating the standard scores of RD_1ij_ and RD_2ij_. In order to show the strength of the association between subject gene and subject entity more intuitively, we sort the standard scores, where a smaller number represents a closer association between subject gene and subject entity. Finally, we obtain the association matrix of all subject genes and subject entities according to the above method.

#### Step 5: Establish association matrix between subject-related antineoplastic drug entities and genes

We have collected the names of antineoplastic drug from multiple data sources, including drug information for 622 antineoplastic drugs included in My Cancer Genome (https://www.mycancergenome.org/)^[10]^, drug information of 267 virtual sieve drugs included in Sanger Drug (https://www.cancerrxgene.org/translation/Drug), and drug information of 62 clinically available antineoplastic drugs (S1 Table) that we have summarized. This list of antineoplastic drugs included kinase inhibitors KI, antibody drugs, antibody-conjugated drugs, chemotherapeutic drugs, hormones, and other classes. We manually performed deduplication and inspections and obtained a total of 803 antineoplastic drug names. We compare the subject-related entity with the name of the antineoplastic drugs in this compiled antineoplastic drug name database and then combine the association matrix between the gene and the subject-related entity obtained from the above process to form the gene-antineoplastic drug association matrix.

### Establish the standard model for antineoplastic drug-X

We selected 18 common antineoplastic drugs to build data interfaces based on the automated literature mining method and separately obtained 18 antineoplastic drug standard models. They were bevacizumab, capecitabine, carboplatin, cetuximab, cisplatin, cyclophosphamide, dexamethasone, doxorubicin, erlotinib, etoposide, fluorouracil, gemcitabine, melphalan, methotrexate, pemetrexed, rapamycin, sorafenib, and vincristine. Each antineoplastic drug-X standard model was constructed as follows: 1) genes with accurate relevance to the antineoplastic drug-X were identified, 2) entities with precise relevance to the antineoplastic drug-X were identified, 3) the association strength between entities and genes was calculated, and 4) relative ranking of the association strengths between antineoplastic drug-X and genes was determined. Finally, for each standard model of antineoplastic drug-X, we obtained the genes and entities precisely related to the antineoplastic drug-X, and obtained the association matrix between antineoplastic drug-X and genes.

### Establish the cancer Y scenario model

Similarly, we selected 16 common cancer types to build data interfaces based on the automated literature mining method and separately obtained 16 cancer scenario models. They were breast cancer, cervical cancer, colon cancer, colorectal cancer, epithelioid sarcoma, esophageal cancer, glioma, hepatocellular carcinoma (HCC), head and neck cancer, Hodgkin lymphoma, melanoma, non-Hodgkin lymphoma, non-small cell lung cancer (NSCLC), ovarian cancer, pancreatic cancer, and thyroid carcinoma. Each scenario model for cancer Y was constructed as follows: 1) genes with accurate relevance to the cancer-Y were identified, 2) entities with precise relevance to the cancer-Y were identified, 3) the association strength between entities and genes was calculated, and 4) alignment with the antineoplastic drug name database was performed to obtain the relative ranking of the association strength between antineoplastic drug entities and genes. Finally, for each cancer-Y scenario model, we obtained the gene entities, antineoplastic drug entities, and the association matrix between antineoplastic drug entity and gene entity that are precisely related to the cancer.

### Model fitting in a binary phase diagram

To evaluate the effectiveness of antineoplastic drugs in different cancer scenarios, we need to obtain the key parameters of antineoplastic drug-X in the standard model and different cancer-Y scenario models, respectively. For the standard model of antineoplastic drug-X, we obtained the number of genes Gx that were precisely related to antineoplastic drug-X and the cumulative association strength (T) between genes and antineoplastic drug-X. For the antineoplastic drug-X in different cancer-Y scenario models, the gene Gy associated with the cancer-Y was obtained and compared with Gx in the antineoplastic drug-X standard model to calculate the number (N) of intersecting genes. Then, we calculated the sum of the association strengths between Gx and antineoplastic drug-X to obtain T based on the association matrix between the gene Gy and antineoplastic drug-X in cancer-Y scenario model. In addition, we need to standardize the association matrix between gene Gy and antineoplastic drug entities to compare the association strength between the same antineoplastic drug-X and gene Gy in different cancer-Y scenarios. In a specific cancer-Y scenario model, we normalized the rank number of the antineoplastic drug-X divided by the number of entities precisely related to cancer-Y. Next, we compared the standard model with the scenario models of various cancers to obtain the scenario model of antineoplastic drug. We placed each antineoplastic drug scenario model based on the two parameters T and N in a binary phase diagram for function fitting and model evaluation. We defined the number N of intersecting genes as the X-axis and the cumulative association strength T as the Y-axis. We fitted the linear regression model according to the number N of the intersecting genes of the antineoplastic drug-X in different cancer-Y scenario models and the cumulative association strength T. Next, the residual value of the model was calculated and the outliers with the minimum negative residual value were eliminated for a better linear fitting to obtain a stochastic scenario model. At the same time, the removed parameter points were fitted to a new linear model to obtain a rational scenario model. Finally, we introduced the standard model of antineoplastic drug into the binary phase diagram: 1) the standard model parameter point was fitted in the rational scenario model parameter points to get the R_1_-square and 2) the standard model parameter point was fitted in the stochastic scenario model parameter points to get the R_2_-square. The optimal linear model was evaluated by comparing R_1_-square with R_2_-square.

### Validation using the Sanger Genomics of drug sensitivity database

We downloaded the publicly available drug sensitivity database from the Genomics of Drug Sensitivity in Cancer (GDSC) (www.cancerrxgene.org), which contains 256 drugs or compounds for more than 1000 tumor cell lines. We combined the drug IC50 data of all tumor cell lines to calculate the average response rate of antineoplastic drugs in tumor cell lines from different sources. To increase the reliability of our results, we selected the stochastic scenario model in the biphasic graph of antineoplastic drug to compare with the tumors enriched with the most genes in the rational scenario model, and then calculated the standard scores of the relative global changes in the average response rate of the antineoplastic drug in the tumor cell lines. For a standard score of Z_stochastic_>Z_rational_, we define the effect of antineoplastic drugs as more sensitive in rational scenario models, whereas the opposite tends to be more resistant effects.

## Results

### Construct gene-knowledge map of antineoplastic drugs in specific cancer scenarios by automated methods

To create the knowledge map in a specific scenario, we automatically build a standard user interface based on genes from a large number of studies in the literature, including the entities and genes associated with the scenario, and the matrix that describes the relationship between the entity and the gene. Using the “ovarian cancer” scenario as an example, our first step was to search PubMed database with the “ovarian cancer” keyword to obtain text data for all peer-reviewed and published in studies relevant to this scenario. On Apr 14, 2018, we had obtained 46,594 articles, including the title of the article, author/unit information, and abstracts. We organized and cleaned the text data and compiled it into the subject dictionary (SD). This dictionary contained entity nouns and gene symbols that were related or unrelated to the scenario, as well as a large number of common entities that were unrelated to the subject scenario. If we directly performed frequency statistics, then the limited sample size results might have significantly reduced interference from noise in the information, but would result in the loss of the valuable associations. In addition, if we used a background text database obtained as a generic control corpus, then we would generate generic entities and gene symbols related to biomedicine, but not necessarily related to the subject scenario. Therefore, to obtain the entities most relevant to the subject scenario, we searched the literature using “public health” as the keyword and compiled this set into the public health dictionary (PD) as the reference corpus, which contains a wide range of commonly used medical related entities and their associations. The comparisons between SD and PD should then exclude specific entities of the non-scenarios and low correlation scenarios (the specific process is shown in Fig 1). In the analysis, we also considered the balance of information by setting the relevant parameters to adjust for the size of the abstract set before carrying out the statistical analyses. Each word in SD was compared with the HUGO Gene Nomenclature Committee (HGNC) database to obtain potential candidate genes with official nomenclature. Common words with the same name as genes were also removed through case verification, such as identifying the difference between the gene ‘WAS’ and the verb ‘was’.

To obtain gene symbols related to the ovarian cancer scenario, we counted the number of abstracts containing each candidate gene in the “ovarian cancer” abstract set and the “public health” abstract set. The gene symbols associated with ovarian cancer scenario were determined by a higher odds ratio (for example, odds ratio⩾6). In this case, we obtained 1,441 gene symbols associated with ovarian cancer. Then, we obtained the subject gene abstract set (SGA) associated with the ovarian cancer scenario based on the distribution of these gene symbols in the text. In this abstract set, we took a similar approach of comparing with the entities in the PD, and finally screened 1,926 biological entities (W1) associated with ovarian cancer, including clinical observations, phenotypes, treatments, drugs, and other related clinical concepts (see Methods). At the same time, entities were rendered case-insensitive, and nouns with plural forms and homologous words with adjectives and adverbs were automatically merged into the same entity and assigned the same number.

After obtaining subject entities and gene symbols associated with the specific cancer scenario, the next step was to build the association matrix between the subject gene and the subject entity. An association is based on whether the gene and the entity co-occur in the same literature abstract. In the ovarian cancer scenario, we mined and linked 1,926 entities with 1,441 genes into a association matrix C (see Methods). Each column in matrix C represents the co-occurrence intensity of a gene with different entities, and the distribution of this intensity may be different for different entities associated with different genes. Therefore, we ranked the associations, with a smaller number representing a stronger association between the gene and the current entity. To extract the antineoplastic drug associated with the specific cancer scenario, we compared the entities in W1 with the antineoplastic drug name database, which gave us an association matrix between 29 antineoplastic drugs and 1,441 subject genes. The approach presented is an automated method to mine and organize relevant knowledge from literature abstracts of specific clinical medical scenarios. This information is provided in the form of a table that is not only convenient for users to read and understand, but also provides standardized input data for future machine learning or artificial intelligence methods.

### Literature mining can verify the recommended drugs in the medication guidelines for ovarian cancer based on a similar mechanism

Based on the abstracts set from published ovarian cancer literature, 29 antineoplastic drugs and 1,441 genes were associated by different intensities. By manually searching the above-mentioned drugs in the antineoplastic drug name database, we confirmed that most of the antineoplastic drugs are widely used in clinical practice for treating ovarian cancer with significant therapeutic effects, such as platinum compounds. We can obtain the gene regulation mechanism via the distribution of the association strength of genes related to different antineoplastic drugs. We screened the genes with relative rankings of the association strengths in the top 10% of all entities as a subset among the 1,441 genes, which we defined as the significantly associated genes of ovarian cancer related to antineoplastic drugs. We further constructed a network based on the association between 29 antineoplastic drugs and significantly associated genes, as shown in Fig 2A. In this network, an association between an antineoplastic drug and a gene is called a linkage. We found that different antineoplastic drugs were usually linked to a group of genes with different intensities (Table 1), and 24 of the 29 antineoplastic drugs had a number of uniquely linked genes that was less than 5% of the total number of genes linked.

**Table 1.** Linkage information between antineoplastic drugs and linked genes in ovarian cancer.

| No | Antineoplastic Drug | The number of genes<br>(relative rank < 10%) | The number of<br>specific genes | The percentage of<br>specific genes | The gene of the<br>most related with<br>drug |
| --- | --- | --- | --- | --- | --- |
| 1 | leucovorin | 500 | 218 | 43.60% | SHE |
| 2 | anastrozole | 282 | 5 | 1.77% | CCND1 |
| 3 | cisplatin | 266 | 10 | 3.76% | ORAI1 |
| 4 | bortezomib | 252 | 8 | 3.17% | ADRM1、UCN |
| 5 | lapatinib | 245 | 2 | 0.81% | MED1、PERP |
| 6 | platinum | 210 | 5 | 2.38% | ERCC1 |
| 7 | erlotinib | 199 | 2 | 1.00% | GALNT6 |
| 8 | gefitinib | 199 | 1 | 0.50% | EMP3 |
| 9 | bevacizumab | 184 | 14 | 7.61% | HIPK3 |
| 10 | herceptin | 169 | 4 | 2.37% | DSG2 |
| 11 | pertuzumab | 163 | 2 | 1.23% | SP3 |
| 12 | sunitinib | 160 | 5 | 3.13% | TEC |
| 13 | pemetrexed | 155 | 1 | 0.65% | ROS1 |
| 14 | everolimus | 152 | 3 | 1.97% | EIF4B |
| 15 | adriamycin | 142 | 4 | 2.82% | HIPK2、IKBKE |
| 16 | doxorubicin | 134 | 6 | 4.48% | TOP1MT |
| 17 | carboplatin | 129 | 2 | 1.55% | TRPM8 |
| 18 | cetuximab | 122 | 2 | 1.64% | CDH2 |
| 19 | vincristine | 104 | 5 | 4.81% | COL12A1、<br>COL1A2、<br>COL21A1、<br>EGFL6、MCL1、<br>POSTN、<br>SLC16A14、<br>SLC2A14 |
| 20 | temsirolimus | 88 | 2 | 2.27% | RHEB |
| 21 | etoposide | 85 | 1 | 1.18% | BCL2L12 |
| 22 | gemcitabine | 76 | 4 | 5.26% | SGK1 |
| 23 | imatinib | 66 | 2 | 3.03% | AMN |
| 24 | anthracyclines | 54 | 5 | 9.26% | CENPF |
| 25 | rapamycin | 53 | 2 | 3.77% | MTOR |
| 26 | sorafenib | 52 | 0 | 0 | PIK3IP1 |
| 27 | cyclophosphamide | 47 | 2 | 4.26% | KLK15 |
| 28 | tamoxifen | 40 | 5 | 12.50% | PTGFR |
| 29 | trastuzumab | 28 | 0 | 0 | ADAM17 |

**Fig. 2.**
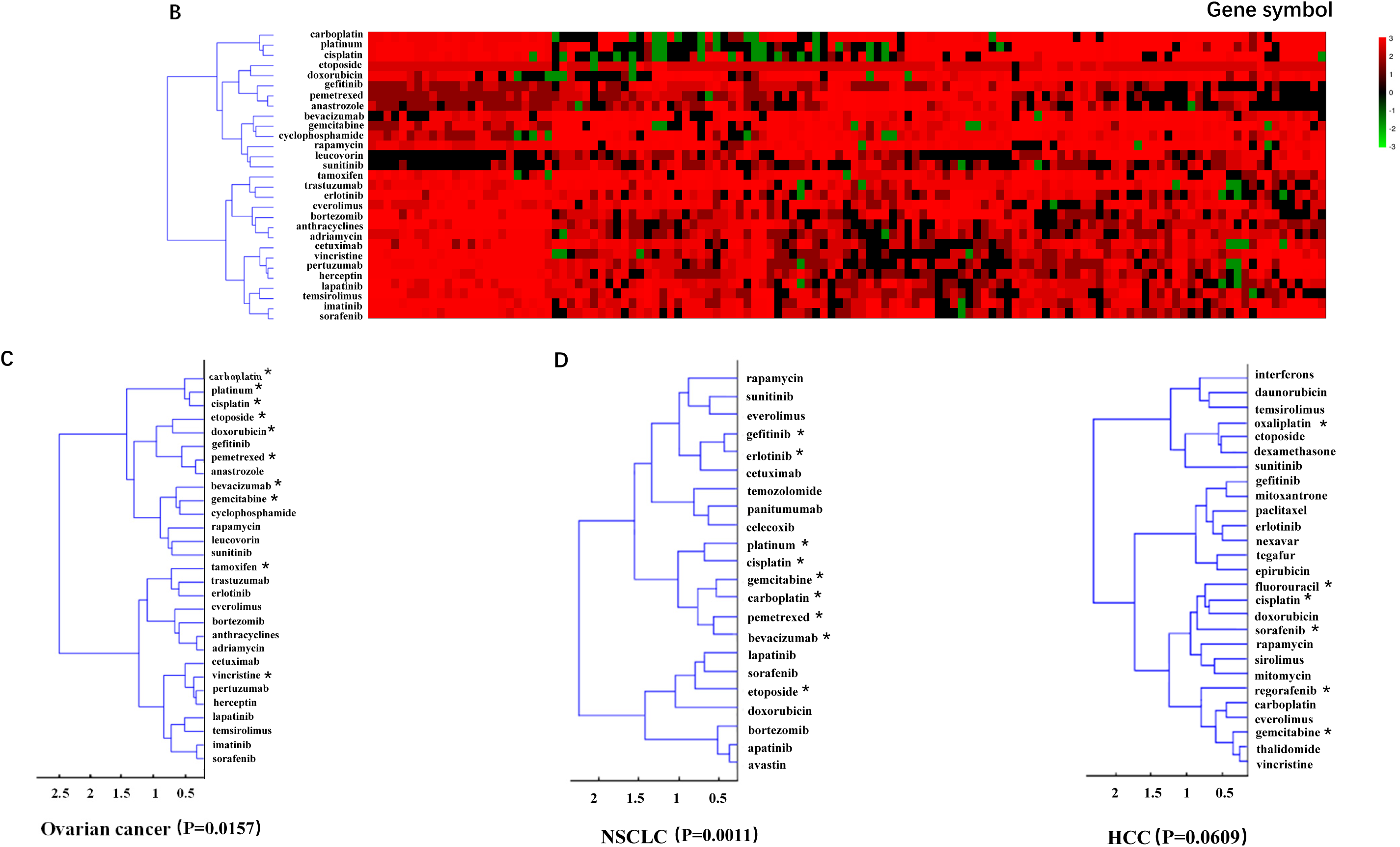
(A) The network analysis of antineoplastic drugs and subject genes in ovarian cancer (relative ranking of antineoplastic drug-gene correlations ≤ top 10%). The edges between them represent the antineoplastic drug-gene interactions. The antineoplastic drugs and the genes considered for the network assembly are highlighted in red and white, respectively. The correlation between antineoplastic drugs and potential genes can be easily observed from this network. (B) Cluster heatmap of the association strengths between 29 antineoplastic drugs and genes in “ovarian cancer” literature abstracts. (C) Clustering map of the association strengths of antineoplastic drugs and genes in ovarian cancer, showing that most NCCN guidelines recommend drugs were significantly clustered into one category (p=0.0157). (D) Clustering map of the association strengths between genes and antineoplastic drugs in non-small cell lung cancer (NSCLC) (P=0.0011) and in hepatocellular carcinoma (HCC) (P=0.0609). The recommend drugs in the NCCN guidelines are labeled by “*”.

The clustering results can more intuitively reflect the strength of a particular research mechanism. Based on the data matrix of antineoplastic drugs and precisely related genes in ovarian cancer scenarios, we performed cluster analysis on the antineoplastic drugs (Fig. 2B). As expected, antineoplastic drugs closer together in such clusters tended to share similar mechanisms of action and were grouped into one category. At the same time, we further explored the correlation between antineoplastic drugs from the perspective of the drugs recommended in clinical practice guidelines. We compared the antineoplastic drugs in the clustering results with the drugs recommended in the National comprehensive cancer network (NCCN) clinical practice guidelines for ovarian cancer. We found 11 drugs were recommended in the ovarian cancer clinical guidelines, including cisplatin, platinum, carboplatin, doxorubicin, gemcitabine, cyclophosphamide, bevacizumab, pemetrexed, tamoxifen, etoposide, and vincristine (labeled with an asterisk in the Fig 2C). Our clustering results showed that most of the antineoplastic drugs recommended by NCCN guideline were significantly clustered into one category (p=0.0359), whereas all antineoplastic drugs were assumed to be classified into two categories (Fig 2C).

We propose two explanations for the above phenomenon:

1. In the ovarian cancer scenario, different antineoplastic drugs may share similar molecular mechanisms, such as platinum-based drugs, cisplatin and carboplatin;
2. Researchers studying the molecular mechanism of antineoplastic drug will begin with a group of commonly used genes according to their expertise.

Different antineoplastic drugs have different mechanisms of action; for example, cyclophosphamide interstrand DNA cross-links to inhibit DNA replication and initiates cell death^[11–13]^, whereas bevacizumab inhibits angiogenic cytokines^[14]^, and similarly, tamoxifen has actions different from platinum compounds^[15, 16]^. However, we found that most of the genes studied in the above drugs were limited to a small number of genes in the large-scale text data. It is possible that a gene will confer different functional meanings in different abstracts in the literature. Therefore, the finding supports the latter of the two previous explanations that there is a preference when studying antineoplastic drug mechanisms, in that researchers tend to scrutinize functions of familiar genes in specific scenarios to explain the possible mechanisms. To test the universality of the above findings, we continued to analyze the intensity distribution of the drug-gene linkage of antineoplastic drugs used in treatments of NSCLC and HCC. The results showed that most of the recommended antineoplastic drugs in NCCN guidelines clustered in the same category with P values ranging from 0.0011 to 0.0609 (Fig 2D), in a similar way to the results for ovarian cancer. These findings show that there is a certain preference in the decision making for clinical treatments for ovarian cancer, lung cancer, liver cancer, and possibly other cancers based on human experience.

The above results raise several questions that need to be addressed. 1) Does the knowledge accumulated by researchers in a specific clinical scenario really follow such a remarkable trend? 2) Is the knowledge in published literature abstracts clinically applicable? 3) Specifically, when an antineoplastic drug is used in the clinical treatment of a specific cancer type, is the basis for the application of such a treatment regimen a rational design or inclined to be a random selection? To address these issues, we need to establish a new quantitative method for rational evaluation.

### Antineoplastic drug-X standard model

In this study, we used a set of genes linked to antineoplastic drugs at different intensities to characterize the extent and depth of the current knowledge of an antineoplastic drug. This set of genes comes from all the knowledge about the antineoplastic drug, i.e., genes associated in different ways with the antineoplastic drug in different clinical or laboratory scenarios. Fig 3 shows the method for obtaining a standard model of an antineoplastic drug. We have obtained standard models for 18 antineoplastic drugs and counted the number of literature abstracts on which each model is based, the number of genes associated with the antineoplastic drug, and the most relevant gene information (Table 2, S2 Table).

**Table 2.** Basic information about Antineoplastic drug standard models.

| No. | Antineoplastic drug | Gene Counts | Antineoplastic drug Counts | the number of Abstracts | The most relevant Gene |
| --- | --- | --- | --- | --- | --- |
| 1 | bevacizumab | 160 | 35 | 15262 | NF2 |
| 2 | capecitabine | 102 | 19 | 5935 | CYP2C9 |
| 3 | carboplatin | 182 | 29 | 15330 | ATP7B |
| 4 | cetuximab | 166 | 18 | 6178 | DDR2 |
| 5 | cisplatin | 848 | 57 | 45053 | LRRC8A |
| 6 | cyclophosphamide | 250 | 47 | 38720 | MICE |
| 7 | dexamethasone | 660 | 41 | 39221 | NAPB |
| 8 | doxorubicin | 641 | 60 | 40304 | CLPTM1' |
| 9 | erlotinib | 203 | 18 | 5696 | CYP2D6 |
| 10 | etoposide | 365 | 38 | 23851 | DTNB |
| 11 | fluorouracil | 396 | 52 | 43583 | ATP5J |
| 12 | gemcitabine | 311 | 35 | 14651 | GEM |
| 13 | melfalan | 96 | 21 | 10362 | BROX |
| 14 | methotrexate | 191 | 41 | 43450 | SP2 |
| 15 | pemetrexed | 92 | 15 | 2869 | SLC22A8 |
| 16 | rapamycin | 927 | 52 | 35931 | MTOR |
| 17 | sorafenib | 233 | 22 | 6968 | FIP1L1 |
| 18 | vincristine | 221 | 31 | 29758 | SARM1 |

**Fig. 3.**
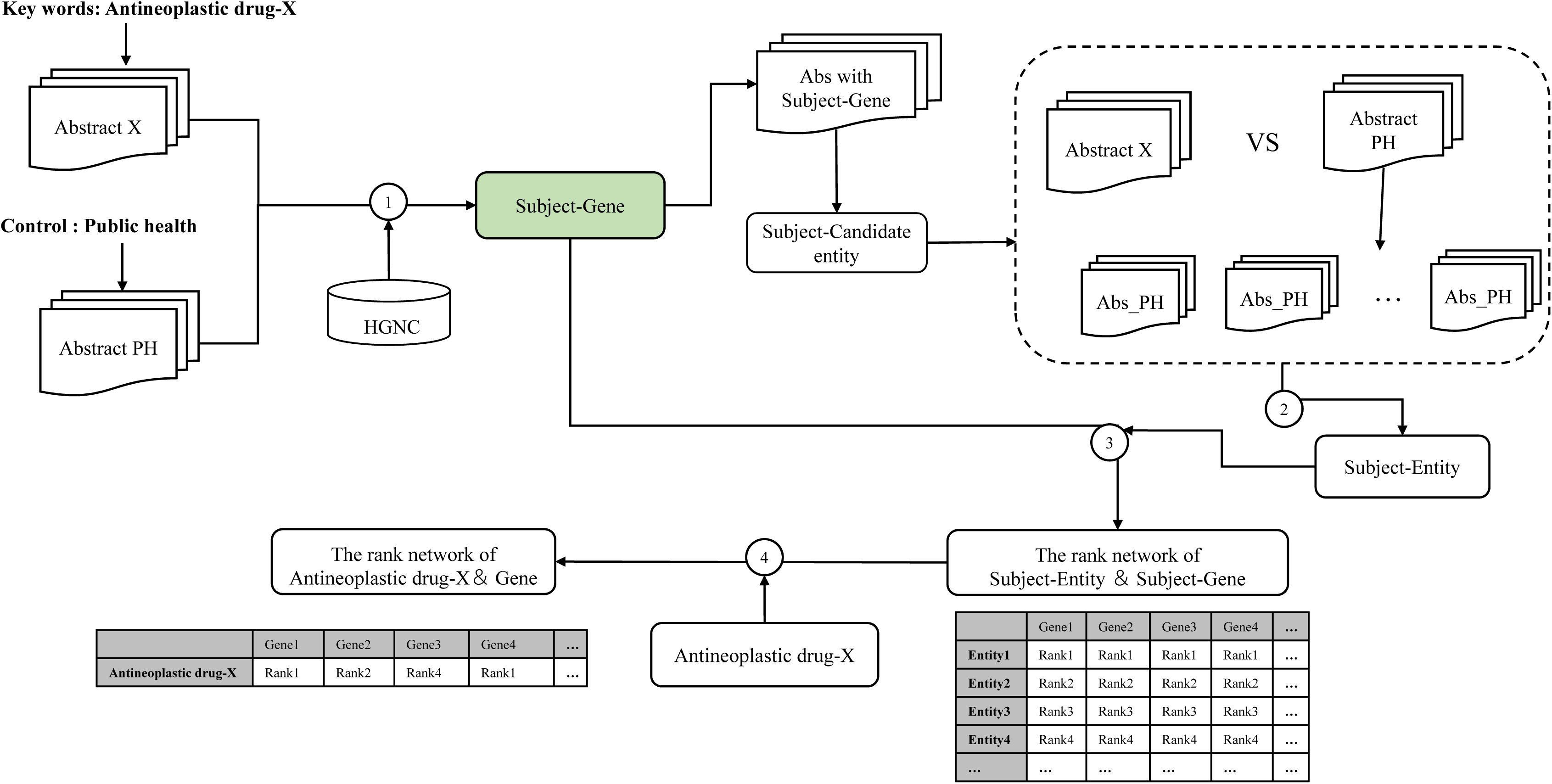
Steps of the antineoplastic drug-X standard model building process.

### Cancer-Y Scenario Model

Similar to the standard model of antineoplastic drugs, we also used a set of genes to describe a cancer that is linked to the related entity at different intensities to reflect the different aspects of the cancer, such as diagnosis, treatment, drug resistance mechanism, or side effects. Fig 4 illustrates the method of obtaining a scenario model for cancer. We have obtained scenario models for 16 cancer types and counted the number of literature abstracts on which each model is based, the number of genes and antineoplastic drugs associated with different cancer types, and the most relevant gene information (Table 3, S3 Table).

**Table 3.** Basic information about cancer scenario models.

| No. | Cancer Type | Gene Counts | Antineoplastic drug Counts | the number of Abstracts | The most relevant Gene |
| --- | --- | --- | --- | --- | --- |
| 1 | Breast cancer | 1777 | 56 | 135179 | BAX |
| 2 | Cervical cancer | 38 | 8 | 3485 | TIMP2 |
| 3 | Colon cancer | 68 | 6 | 3761 | BRAF |
| 4 | Colorectal cancer | 103 | 15 | 10215 | NRP1 |
| 5 | Epithelioid sarcoma | 985 | 32 | 41893 | MTOR |
| 6 | Esophageal cancer | 598 | 12 | 56385 | PIK3CA |
| 7 | Glioma | 1154 | 35 | 80950 | ASB1 |
| 8 | Head and neck cancer | 77 | 14 | 9427 | BLM |
| 9 | Hepatocellular Carcinoma (HCC) | 1328 | 40 | 94395 | ARF6 |
| 10 | Hodgkin lymphoma (HL) | 157 | 15 | 31503 | EFS |
| 11 | Melanoma | 560 | 78 | 59508 | PTN |
| 12 | Non-hodgkin lymphoma (NHL) | 151 | 18 | 18546 | GAN |
| 13 | Non-small cell lung cancer (NSCLC) | 1098 | 27 | 35523 | ERBB3 |
| 14 | Ovarian cancer | 1441 | 29 | 42441 | MTOR |
| 15 | Pancreatic cancer | 1088 | 36 | 88273 | BID |
| 16 | Thyroid carcinoma | 516 | 13 | 65515 | TSN |

**Fig. 4.**
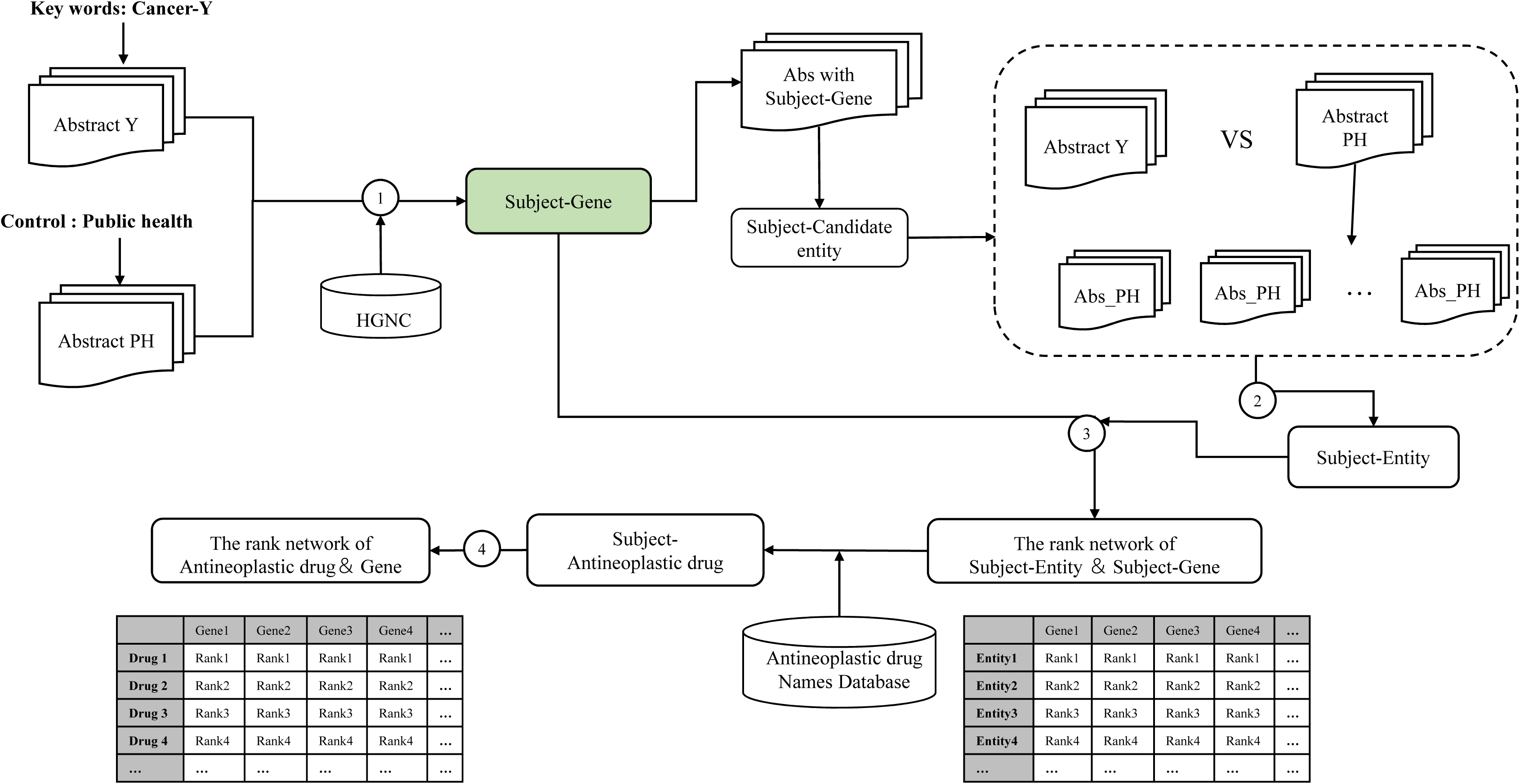
Steps of the cancer-Y scenario model building process.

### Non-empirical dependent assessment of an antineoplastic drug used in a specific cancer Binary phase diagram

The standard model of antineoplastic drug-X shows gene Gx associated with an antineoplastic drug. Two parameters of Gx can be observed in the scenario model of cancer-Y: 1) the cumulative association strength T of Gx with antineoplastic drug-X in the cancer-Y scenario model, and 2) the number of intersecting genes N of Gx in the cancer-Y scenario model. The former shows that the antineoplastic drug-X in cancer-Y scenario has a stronger association with related genes suggesting that the mechanism of the antineoplastic drug may be studied more frequently in this cancer-Y scenario (emphasizing the depth of research), whereas the latter reflects the degree of overlap between the study of the mechanisms of antineoplastic drug-X and cancer-Y. This is related to the degree of research on antineoplastic drug-X itself, as well as on the degree of research on antineoplastic drug-X in cancer-Y scenario (emphasizing the breadth of research). The specific calculation methods of T and N can be found in Methods. Therefore, we represent these two parameters on a binary phase diagram and obtain the scenario models of antineoplastic drug according to T and N parameters of antineoplastic drug-X in different cancer-Y scenarios.

### Stochastic model

In the stochastic model, we compare the standard model of antineoplastic drug with the scenario models of multiple cancers to obtain various scenario models of the antineoplastic drug, and then mark the parameters T and N in each antineoplastic drug scenario model on a binary phase diagram. For example, with methotrexate, we can clearly observe that each point on the binary phase diagram falls near a straight line, reflecting that there may be a positive proportional relationship between T and N. Therefore, we fit the function with the parameters T and N, which is equivalent to random sampling in the standard model of antineoplastic drugs with the number of intersecting genes N of different antineoplastic drug scenario models as variables, and calculate the corresponding cumulative association strength T to fit the function. The results showed a linear function, indicating that the cumulative association strength increases proportionally with the increase in the number of intersecting genes N. Furthermore, when linearly fitting each point of the above different antineoplastic drug scenario models, we found the standard model of antineoplastic drugs fell near the extended line, indicating that the linear positive correlation actually reflects a randomness. Therefore, we call this model a stochastic model (Fig 5).

**Fig. 5.**
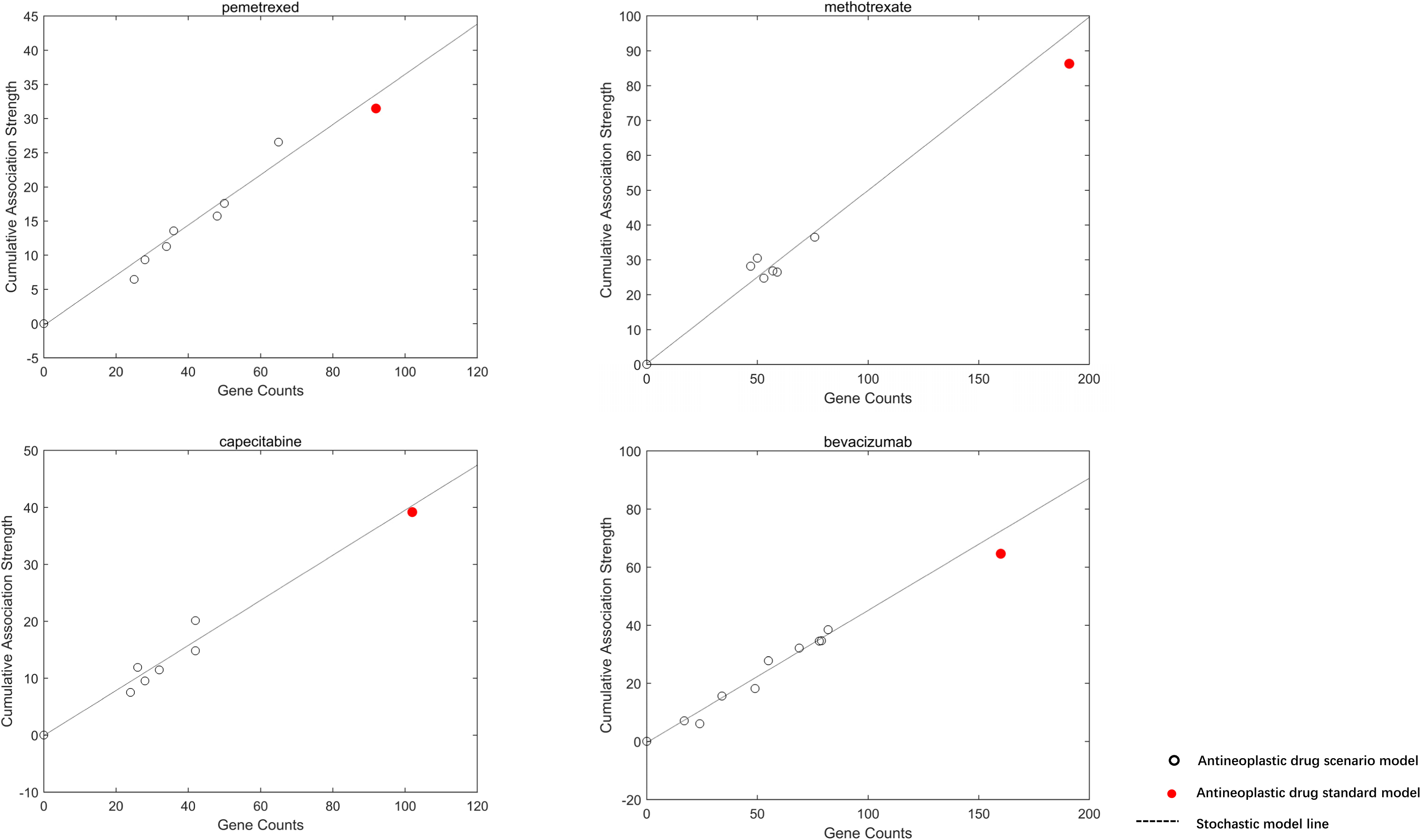
The scatter plot of the standard model and multiple scenario models of antineoplastic drugs. The linear regression line (stochastic model line) shown on the scenario models illustrates the similar distribution trends between the standard model and multiple scenario models of antineoplastic drug.

### Non-empirical independent model

However, we found that the parameter points obtained by certain antineoplastic drugs in different cancer scenarios were not a perfectly random distribution, such as cisplatin. Therefore, we adopted an algorithm that gradually eliminates outliers for a better linear fitting by removing some parameter points to obtain the linear stochastic scenario model. With a good enough fitting model, we found that the excluded parameter points could also fit a new linear model called a rational scenario model (Fig 6). The results showed a variety of antineoplastic drugs displayed this characteristic, suggesting some antineoplastic drugs may have substantially different mechanisms in different cancer scenarios.

**Fig. 6.**
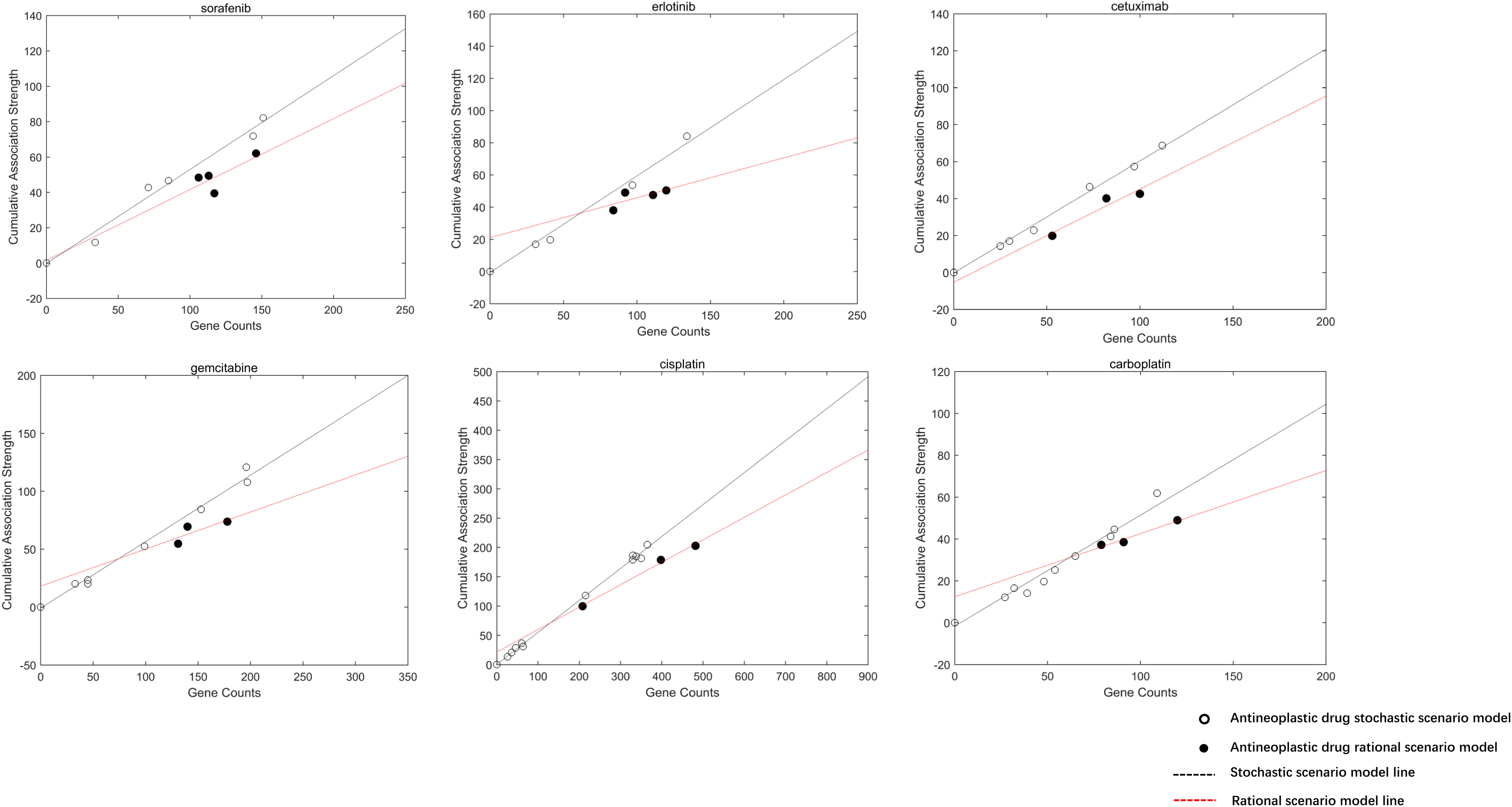
The scatter plot of multiple scenario models of antineoplastic drugs. The distribution trend of rational scenario model and random scenario model by fitting the linear regression line to the parameters of the rational scenario model (rational scenario model line) and the parameters of the stochastic scenario model (stochastic scenario model line).

In the binary phase diagram of antineoplastic drugs, we found a characteristic of the rational scenario model in that when the number of genes was the same, the cumulative correlation intensity T was smaller than in the stochastic scenario model, suggesting that a cancer represented by the parameter points in rational scenario model might actually be more closely related to the current antineoplastic drug. To verify this, we added the standard model of the antineoplastic drug into the binary phase diagram together with rational scenario model. We found the linear model fitted by the parameter points of the standard model and rational scenario model was better than the linear model fitted by the parameter points of the standard model and stochastic scenario model (Fig 7). From the above findings, we can infer that the clinical application of antineoplastic drugs such as cisplatin and erlotinib are recommended for ovarian cancer based on research evidence. Whereas gemcitabine in NSCLC may have a tendency to be recommend based on human experience, indicating the research on mechanism of antineoplastic drugs is less applicable across different cancers and that there exists a possible disconnect between basic research and clinical application.

**Fig. 7.**
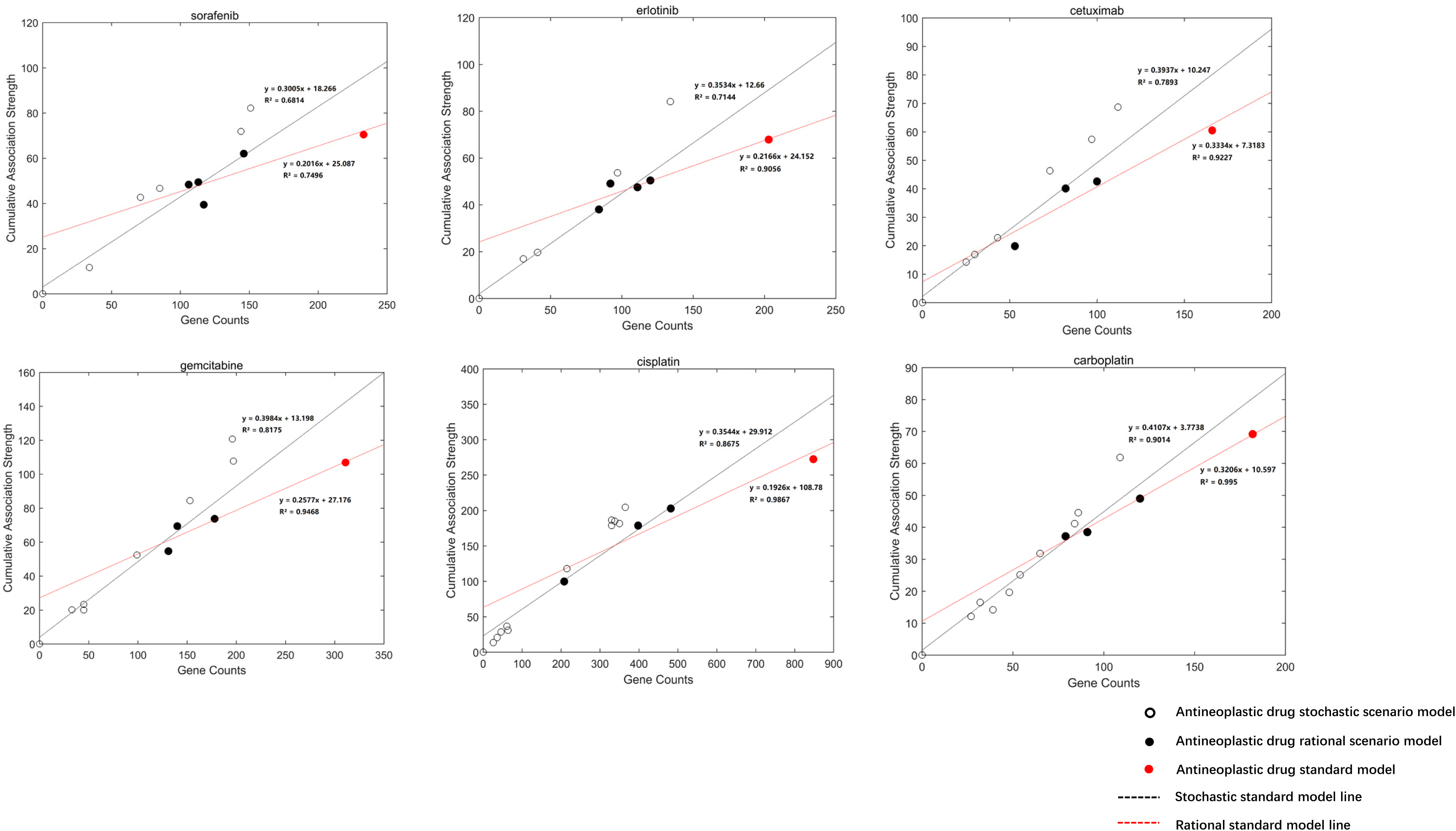
The scatter plot of the standard model and multiple scenario models of antineoplastic drug with linear regression lines. The rational linear regression model was fitted with the parameters from rational scenario models and standard model of antineoplastic drugs; the simulated linear equation, R^2^ value, and the rational standard model line are shown. At the same time, the stochastic linear regression model was fitted with the parameters of the stochastic scenario model and the standard model of antineoplastic drugs; the simulated linear equation, R^2^ value, and the stochastic standard model are shown.

The clinical application of antineoplastic drugs guided by research, in theory, should be more effective. We analyzed the pharmacodynamic data of antineoplastic drugs in different cancer types using the Genomics of Drug Sensitivity in Cancer (GDSC) database to obtain the average sensitivity of antineoplastic drugs to tumors containing the largest number of genes in the stochastic scenario model and rational scenario model, respectively. We found the response rate of tumor cells to cisplatin, gemcitabine, erlotinib, and sorafenib on a stochastic scenario model was lower than that of tumor cells on a rational scenario model (Z_stochastic_>Z_rational_, Fig 8). This suggests the guidelines are actually a hybrid system based on human experience and research knowledge that may not be the most effective for specific application scenarios.

**Fig. 8.**
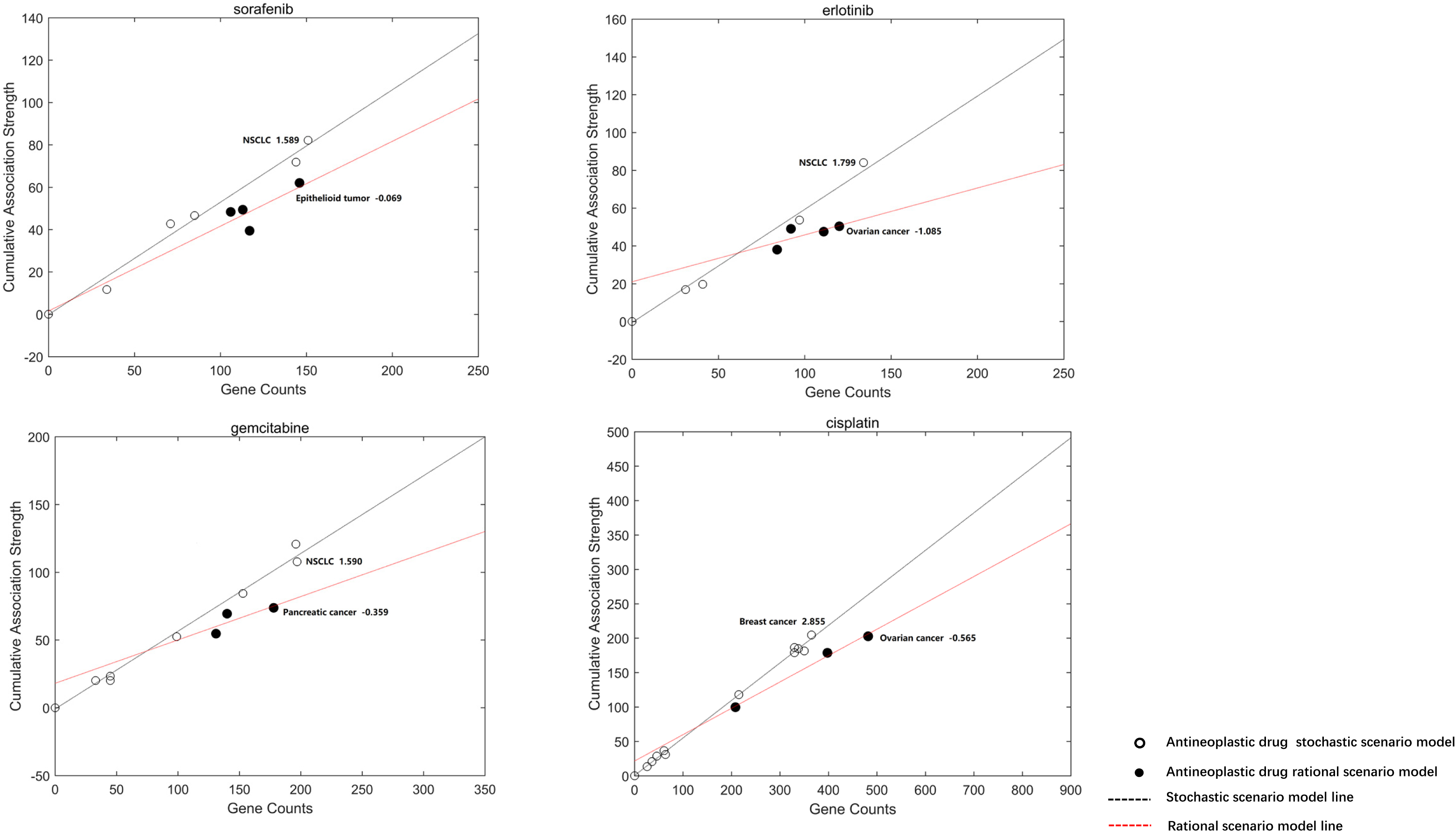
Multiple linear regression models of four antineoplastic drugs in different cancer types in the binary phase diagram. The standard scores are the relative global changes in the average response rate of antineoplastic drugs in the cancers containing the largest number of genes in stochastic and rational scenario models, respectively.

## Discussion

Collating and mining information from the literature has become important approaches for biological knowledge discovery and biomedical research. The biomedical literature is growing exponentially and abstracts contain a large number of experimental results, gene-phenotype description, and pharmacodynamic information. Currently, most biomedical literature mining research related to drugs focus on several aspects. 1) Functional information of genes, such as building structured resources for drug-gene correlation, providing intuitive graphical user interfaces and documented application programming to query the correlations between the gene and the drug^[17–24]^. 2) Identifying molecular biomarkers of drug efficacy in cancer patients and providing evidence for precision medicine by extracting drug-gene correlation information from published literatures, databases and other web resources^[25–28]^. 3) Using machine learning to identify the most effective pharmacogenomic information for drug repositioning^[29–35]^. 4) Predicting drug side effects^[36–38]^, susceptibility, and antitumor drug resistance, to guide hypothesis-driven basic scientific research^[39]^.

Nonetheless, the purpose of our research is different from the previous studies, in that the main aim is to evaluate the effectiveness of common antineoplastic drugs in different clinical scenarios of cancer, particularly from the view of drugs recommended by clinical guidelines. Our definition of effectiveness in this context is that the more studies in the literature that report the molecular mechanisms of antineoplastic drugs on a certain cancer, the higher the probability that the mechanism has being fully elaborated, and thus the antineoplastic drug would be more effective as a clinical treatment for the cancer, which would facilitate the decision to use it. Therefore, we quantitatively determined the association between antineoplastic drugs and genes based on the literature abstracts of studies on specific cancer scenarios using our automated literature mining method. The dataset was displayed in a format that users could understand. We obtained 18 antineoplastic drug standard models and 16 cancer scenario models. The confidence evaluation parameters of these models were extracted and fitted to the multiple linear regression model. We found six antineoplastic drugs were effective in some cancer scenarios, which were validated using high-throughput antineoplastic-drug screening database, such as the GDSC database.

To increase the accuracy of genes associated with antineoplastic drugs in the automated literature mining methods, we used “public health” literature abstracts containing rich entities as the random corpus for the comparison study. Public health covers a wide variety of disciplines ranging from social sciences to business to biological sciences, such as flu pandemics, disaster preparedness, and obesity. Therefore, the probability of each disease or gene being mentioned in public health-related literature should be the same, which excludes specific entity information in non-scenarios and low-association scenarios. Meanwhile, we used “gene” as the prerequisite to identify all the entities in the literature abstracts that were accurately related to genes with high co-occurrence frequency, and then quantified the degree of association between the gene and the entity. This process increases the comprehensiveness of the description of the subject and its association with genes. To prove that the recommended drugs in the clinical guidelines for ovarian cancer, such as the NCCN clinical practice guidelines, are based on similar research mechanisms, we conducted cluster analyses based on the distribution of the association strengths between genes and antineoplastic drugs. The antineoplastic drugs recommended by NCCN for a specific scenario were significantly clustered into one category. For example, carboplatin has a close relationship with cisplatin and are both clustered into one category. We found that ABCC3, TRPM8^[40, 41]^, ATOX1^[42, 43]^ genes were closely related to carboplatin and cisplatin by querying the association matrix between genes and antineoplastic drugs. For example, ABCC3 is a member of ATP binding cassette (ABC) transporter family. Carboplatin chemotherapy induces hyaluronan production which can contribute to chemoresistance by regulating ABC transporter expression^[44]^. A study reported that the resistance gene ABCC3 was co-expressed with lncRNA CTD-2589M5.4 by integrating the published data with data on cisplatin resistant lncRNA in ovarian cancer cell lines or ovarian cancer patients^[45]^. There appears to be a tendency to study antineoplastic drug mechanisms in terms of researcher experience of familiar genes in specific scenarios, which when referenced multiple times, reinforces these possible mechanisms.

Finally, to evaluate the effectiveness of the 18 antineoplastic drugs in 16 cancer scenarios, we performed the literature mining analysis of 16 cancer types and 18 antineoplastic drugs to generate 34 data interfaces. The data interface for the antineoplastic drug was used as a standard model to describe the degree of current understanding of the antineoplastic drug at the gene level. Similar to the standard model of the antineoplastic drug, we also used a set of genes to describe a cancer type and associated the genes with the cancer-related entities at different intensities, and generated data interfaces for different cancer types as the cancer scenario models. In order to evaluate the effectiveness of antineoplastic drugs in different cancer scenarios, we compared the standard model of antineoplastic drug with a variety of cancer scenario models to obtain the cumulative association strength T of gene Gx with antineoplastic drug-X in cancer-Y scenario model, and the number of intersecting genes N of Gx in cancer-Y scenario model. The above two parameters were placed on a binary phase diagram to fit the antineoplastic drug scenario model. The model is equivalent to random sampling in the standard model of antineoplastic drug. By taking the number of intersecting genes N as a variable and calculating the cumulative association intensity corresponding to each variable, the number of intersecting genes N and the cumulative association strength T of antineoplastic drugs was found to be a linear function of N. This indicates that there is a positive quantitative relationship between T and N in the binary phase diagram, and the cumulative association strength increases proportionally with the increase in the number of genes. We obtained a linear fitting of the parameter points representing different antineoplastic drug scenario models. When the standard model of antineoplastic drug falls near the fitted line, we called it a stochastic model. When the standard model of antineoplastic drug falls below the fitted line, we use this standard model as a reference point that shows some antineoplastic drug scenario models could be better linearly fitted with this reference point, which we then called a rational scenario model. As X-axis represents the number of genes and the Y-axis represents the cumulative association strength between genes and antineoplastic drugs, the cumulative association strength of genes and antineoplastic drugs thus deviate from the random distribution in the specific cancer scenario. At the same time, the cumulative association strength decreases, indicating that the functional association between genes and antineoplastic drugs may be closer in this cancer scenario.

Therefore, the validity of antineoplastic drugs in different cancer scenarios can be well evaluated by model fitting in the binary phase diagram. In addition, we used the antineoplastic drug-relative sensitivity data of more than 1000 tumor cell lines in the Genomics of Drug Sensitivity in Cancer (GDSC) database for the verification. It was found that the analyzed tumor cell lines were significantly more sensitive to the antineoplastic drugs used in rational scenario models than in stochastic scenario models. For example, we found that the sensitivity of cisplatin in ovarian cancer was higher than that in breast cancer. A variety of clinical studies have reported that cisplatin combination therapy can be used for the effective treatment of ovarian cancer^[46–50]^. Ovarian cancer was highly sensitive to this chemotherapy compared to many other types of cancer as shown by the overall 5-year survival of over 50%^[51]^. Similarly, cisplatin has been widely used for the treatment of patients with breast cancer ^[52–54]^, with efficiency of only 25% but its effectiveness is still unclear. Cisplatin chemotherapy has high activity in women with a BRCA1 mutation and metastatic breast cancer, with a complete remission rate of 61%^[5555]^. Based on the above results, our method was shown to be a valid and effective tool to evaluate the rationality of medicine decision depending on the conformity of pharmacological mechanisms in the research and in a clinical setting.

## Conclusions

Our literature mining method provides a practical tool to evaluate the applicability of an antineoplastic drug in various cancer types. This study combined automated knowledge-driven methods to establish the antineoplastic drug standard model and cancer scenario models based on the accurate antineoplastic drug-gene association matrix for global or specific cancer scenarios. Then, we used a linear regression analysis method based on the parameters in above models to determine the possible efficacy of an antineoplastic drug in different cancer types. The results can be verified by the Genomics of Drug Sensitivity in Cancer (GDSC) database. This approach has two advantages: 1) it assesses the efficacy of antineoplastic drug in various cancer types to allows more accurate judgments of the use of antineoplastic drugs in clinical practice providing better clinical benefits to patients, and 2) it is a general method that provides a comparable quantitative description of the association between generic entities and genes to delineate molecular mechanisms, and can present a comprehensive knowledge landscape in specific research scenarios for researchers in a specific research field.

## Supporting information

Information of 622 antineoplastic drugs.

The association matrix between antineoplastic drugs and genes in 18 antineoplastic drugs

The association matrix between antineoplastic drugs and genes in 16 cancer types.

## Abbreviations

GDSC: Genomics of Drug Sensitivity in Cancer
HCC: Hepatocellular carcinoma
HBV: Hepatitis B virus
SD: Subject dictionary
PD: Public health dictionary
HGNC: Hugo gene nomenclature commission
SGA: Subject gene abstract set
AGi: Subject gene abstract set containing gene Gi
NAGi: Subject gene abstract set that does not contain gene Gi
SAGi: Random subject gene abstract set containing gene Gi
SNAGi: Random subject gene abstract set that does not contain gene Gi
SPE: Subject-specific entity dictionary
SHE: Subject-shared entity dictionary
HEj: Subject-shared entity j
PEk: Subject-specific entity k
RD: Relationship distance
NSCLC: Non-small cell lung cancer
NCCN: National comprehensive cancer network
ABCC3: ATP Binding Cassette Subfamily C Member 3
TRPM8: Transient Receptor Potential Cation Channel Subfamily M Member 8
ATOX1: Antioxidant 1 Copper Chaperone
ABC: ATP binding cassette
BRCA1: BRCA1 DNA Repair Associated
IC50: Half maximal inhibitory concentration;

## Declarations

### Consent for publication

Not applicable.

### Ethics approval and consent to participate

Not applicable.

### Availability of data and material

All data generated or analysed during this study are included in this published article (Supplementary file 2 and Supplementary file 3).

### Competing interests

The authors declare that they have no competing interests.

### Funding

This work was supported by the National Natural Science Foundation of China (grant no. 31360547).

### Authors’ contributions

GN and XJ conceived and designed the experiments. GN and XJ performed the experiments and analysed the data. GN, XJ, RN and ZF wrote the paper. ZZ set up the website. All authors discussed the results and contributed to the final manuscript.

---

Supplementary file 1.

Information of 622 antineoplastic drugs.

Supplementary file 2.

The association matrix between antineoplastic drugs and genes in 18 antineoplastic drugs.

Supplementary file 3.

The association matrix between antineoplastic drugs and genes in 16 cancer types.

